# When enemies do not become friends: Experimental evolution of heat-stress adaptation in a vertically transmitted parasite

**DOI:** 10.1101/2020.01.23.917773

**Authors:** Eike Dusi, Sascha Krenek, Thomas Petzoldt, Oliver Kaltz, Thomas U. Berendonk

**Author notes:** Author contributions: ED, SK and TUB conceived the study; ED conducted the experiment. OK, ED and TP conducted the statistical analyses. OK and ED wrote the manuscript; SK, TP and TUB provided comments on the manuscript. All authors gave final approval for submission.

## Abstract

Strict vertical (parent-to-offspring) transmission of symbionts may be associated with benevolence and mutualism, because host reproduction and symbiont transmission are linked. Nonetheless, many such symbionts do reduce host fitness costs and the factors driving their evolution are unclear. Using microcosm populations of *Caedibacter taeniospiralis*, a vertically transmitted bacterial symbiont of the protozoan *Paramecium tetraurelia*, we investigated evolutionary change under permissive conditions (26°C) and under heat stress (32°C), over 150 host generations. We found evidence for symbiont adaptation to heat stress, involving increased infection persistence and, consequently, higher rates of vertical transmission. For certain genetic backgrounds, heat-stress adaptation was associated with higher proliferation of infected lines at 26°C, suggesting cost-free evolution of super-generalists. Despite a general decrease in symbiont virulence relative to ancestral strains, infected evolved lines still showed substantial reductions (>20%) in population growth. There was no sign of a temperature-specific reduction in virulence or protection of the host against heat stress. Our results show that obligately vertically transmitted symbionts can still qualify as ‘parasites’ and evolve adaptations that do not help their hosts. Long-term infection persistence might require additional mechanisms, such as the symbiont-mediated ’killer trait’ in this system, allowing the selective elimination of uninfected individuals in the population.

## Introduction

Symbiosis generally involves transfer of resources and services between the two interacting species, thereby generating reciprocal impacts on their fitness (Thrall et al. 2007; Leung and Poulin 2008). The outcome of an interaction, from positive (mutualism) to neutral (commensalism) to negative/antagonistic (parasitism), depends on the net effect of costs and benefits for the players involved. Over the past years, an increasing number of studies have suggested that this balance between costs and benefits is strongly condition-dependent, with systems potentially shifting back and forth on a mutualism-parasitism continuum (Michalakis et al. 1992; Brown 2003; Restif and Kaltz 2006; Thompson and Fernandez 2006; Wolinska and King 2009; Vale et al. 2011). Factors determining sign and strength of an interaction are the symbiont transmission mode, environmental conditions, as well as the genetic background of symbiont and host (Ewald 1987; Wolinska and King 2009). Most empirical and experimental work has highlighted the short-term consequences of variation in these factors over one or very few generations (Thomas and Blanford 2003; Wolinska and King 2009; Ebert 2013), but it is still less clear how they drive long-term evolutionary and co-evolutionary processes (Mahmud et al. 2017; Shapiro and Turner 2018; Hatcher et al. 2005).

Symbionts with exclusive vertical transmission are particularly interesting in this context. Vertical transmission occurs from infected parents to offspring, and therefore symbiont transmission success is directly linked with host reproduction (Fine 1975). Unlike in systems with infectious horizontal transmission, allowing some degree of host exploitation and damage (Alizon et al. 2009), symbiont and host fitness are positively aligned, so that vertically transmitted symbionts should evolve to avoid harm to their host or even become beneficial (Ewald 1987; Jones et al. 2007; Ebert 2013). Moreover, under exclusive vertical transmission, the symbiont is locked up in a single line of descendant. This may lead to the accumulation of deleterious mutations or loss of function (Dale and Moran 2006; Feldhaar 2011), but also facilitate specialisation and co-evolution (Brucker and Bordenstein 2012; Jaenike 2015). These ideas are consistent with the observation that some of the major evolutionary transitions from parasitic to mutualistic relationships are associated with a switch from horizontal to vertical transmission, as shown across the phylogenetic tree (Moran et al. 2008; Sachs et al. 2011), but also in microcosm experiments (e.g., Dusi et al. 2015; Shapiro and Turner 2018).

Nonetheless, many vertically transmitted symbionts are known to be harmful to their host, qualifying them as parasites (Kelly et al. 2003; Mouton et al. 2004; Ebert 2013; Dusi et al. 2014). In some cases, this can be explained by the existence of residual horizontal transmission, offsetting the negative effects on host fitness. In other cases, symbionts manipulate host’s reproductive system in such a way that uninfected individuals are eliminated from the population (e.g., cytoplasmic incompatibility or male killing in *Wolbachia*, thereby increasing the frequency of the symbiont in the population despite its actual fitness costs (Werren 1997; Dunn and Smith 2001; Dunn et al. 2001). Alternatively, vertically transmitted symbionts may provide benefits against natural enemies (Oliver et al. 2005; Haine 2008; Brownlie and Johnson 2009; Jones et al. 2011), competitors or adverse environmental conditions, such as pollutants or high-temperature stress (Russell and Moran 2006; Douglas 1998). Such condition-dependent benefits may open the evolutionary avenue towards mutualism (Fellous and Salvaudon 2009).

We still know very little about how environmental variation impacts the evolutionary dynamics of systems with obligate vertical transmission. For example, in *Buchnera*, a bacterial symbiont of aphids, a single point mutation in the symbiont genome seems to drive a balanced polymorphism of wild-type strains that confer heat-stress protection, but impose fitness costs at lower temperature, and non-protecting mutants that bear no fitness costs at lower temperatures (Dunbar et al. 2007). Variation in heat-stress protection also exists among genotypes (or species) of other aphid-symbionts, indicating a genetic basis on which selection can act (Russell and Moran 2006; Cayetano and Vorburger 2013). Similarly, genetic variation in temperature sensitivity is known for within-host density of *Wolbachia* spp., a widely distributed bacterial symbiont of arthropods (Mouton et al. 2003), although it is less clear how this effect impacts host fitness and thus potential responses to selection (Mouton et al. 2007).

Typically, the above-mentioned experimental work on temperature effects is based on single host individuals and their offspring, spanning little more than one host generation and thereby limiting information on selective, population-wide processes (but see Rouchet and Vorburger 2014). In the present study, we investigated how heat stress conditions affect evolution in experimental microcosm populations of the freshwater protozoan *Paramecium tetraurelia,* infected with the strictly vertically transmitted bacterium *Caedibacter taeniospiralis*. Vertical transmission occurs through the segregation of bacterial cells into the daughter cells of the asexually dividing host. Infection with the symbiont reduces host fitness (Dusi et al. 2014; but see Grosser et al. 2018), but it also confers a so-called ’killer trait’ to the host, leading to the selective killing of uninfected conspecifics in the population (Sonneborn 1943; Schrallhammer and Schweikert 2009). This effect is similar to the male-killing or cytoplasmatic incompatibility of *Wolbachia* or *Cardinium*, which also promotes an increase in the frequency of symbiont carriers in the population by eliminating non-carriers (Hurst et al. 2000; Gotoh et al. 2007).

In a long-term experiment, we exposed infected populations with 5 different genotype associations to a 32°C high-temperature treatment and to a 26°C control treatment. In a previous short-term experiment, infection prevalence had declined at 32°C, indicating limited heat tolerance of the symbiont (Dusi et al. 2014). Thus, the first objective of this study was to test whether this system can evolve in such a way that infection is maintained at the stressful temperature (32°C). Moreover, Grosser et al (2018) demonstrated that *Caedibacter* infection is associated with increased expression of Heat Shock Protein (HSP70), a protective agent against temperature stress (Sorensen et al. 2003). Our second objective was therefore to test whether adaptation to high temperature involved the evolution of reduced virulence or even beneficial effects of infection on its host. After ca. 150 host generations, we performed adaptation assays, comparing ancestral and evolved host-symbiont lines for their capacity to maintain infection and their growth performance. Lines were assayed at both temperatures (26°C and 32°C) to test whether adaptation to the high-temperature environment traded off with the performance at the original permissive temperature.

## Methods

### STUDY SYSTEM

The gammaproteobacterium *Caedibacter taeniospiralis* (Preer and Preer 1982) occupies the cytoplasm of its host *Paramecium tetraurelia,* a cosmopolitan fresh-water ciliate (Ciliophora). Transmission occurs vertically during mitotic host cell division, from the infected mother cell to the two resulting daughter cells. Free-living stages or horizontal transmission are unknown; quasi-horizontal transmission during sexual conjugation is theoretically possible, but has to our knowledge never been documented. Infection reduces host fitness (Dusi et al. 2014), but the parasite also confers a so-called ’killer trait’ to the host, leading to the selective elimination of uninfected *Paramecium* in the population (Schrallhammer and Schweikert 2009). Infection prevalence is stably maintained at temperatures ≤26°C, but rapidly declines at 32°C (Dusi et al. 2014). Thus high temperature cures *Paramecium* from infection; it also suggests that the killer-trait is inefficient at this temperature, as cured cells would otherwise be killed by infected cells (Grosser et al. 2018).

### LONG-TERM SELECTION EXPERIMENT

The long-term selection experiment employed five *Paramecium tetraurelia* strains, each associated with its own *Caedibacter taeniospiralis* symbiont (Preer et al. 1974). Stock cultures of these strains are maintained in 0.25% Cerophyl medium (Krenek et al. 2011), inoculated with *Raoultella planticola* (DMSZ 3069) at 22°C.

Following the results from Dusi et al. (2014), we established a stressful high-temperature selection treatment at 32°C, and a permissive control treatment at 26°C, close to the optimum for *Paramecium* growth and allowing stable persistence of *Caedibacter*. For each strain, three long-term replicate populations (referred to as ’selection line’, hereafter) were assigned to each selection temperature, giving a total of 30 populations (5 host strains x 3 replicates x 2 selection temperatures; Fig. S1). Selection lines had an initial host cell density of 500 cells ml^-1^ in 40ml of culture medium (60ml flasks); initial infection prevalence (i.e., the proportion of infected hosts) was 100%. The selection lines were kept in computer-controlled water baths (for details, see Dusi et al. 2014). Several water baths were assigned to each of the two treatment temperatures, and cultures randomly assigned to a given water bath. Over the course of the experiment, we regularly changed the position of the cultures within and between water baths of the same temperature. Every 2-3 days, 20ml of culture were replaced with freshly bacterized medium. This ensured regular periods of population growth, with approximately three host generations per week. We measured host density and infection prevalence in 2-month intervals (approximately every 25 host generations), as described in Dusi et al. (2014). Note that prevalence data for individual selection lines are only available for the second half of the experiment.

Ancestral cultures were maintained at 10°C, with 25% of the population replaced with freshly bacterized medium every three weeks. This protocol minimised cell division (≈ 1 complete population turnover in 12 weeks) over the 52 weeks of the long-term experiment.

### ADAPTATION ASSAY

After 52 weeks (ca. 150 host generations), we performed an adaptation assay (Fig. S1). Three single *Paramecium* cells were isolated from each selection line, washed in sterile Dryl’s medium (Dryl 1959) and then grown up individually as monoclonal lines for 8 days at the temperature they had experienced during the long-term experiment (26°C or 32°C). We also established monoclonal lines from the ancestral culture of each strain, kept at 10°C during the long-term experiment and re-acclimatised for 8 days at 22°C prior to use. Next, each monoclonal line was split and one half was cured from infection by adding the antibiotic streptomycin (Krenek et al. 2012; Dusi et al. 2014). The presence of infection and its absence after antibiotic treatment was verified using FISH and PCR techniques (Dusi et al. 2014).

Infected and cured monoclonal lines were tested at both 26°C and 32°C. To this end, each monoclonal line was further split into two pre-cultures (Fig. S1). One remained at its original selection temperature; the other was acclimatised in steps of +/- 2°C per day to the ’foreign’ assay temperature and remained at this temperature for another two days prior to the assay. In the same way, ancestral monoclonal cultures were acclimatised to the two temperatures.

From the acclimatised pre-cultures, we started individual assay replicates in flasks containing 20 ml of culture medium, at initial densities of ca. 25 cells ml^-1^ and 100% infection or 0% infection (cured replicates). The flasks were placed in water baths at assay temperatures of 26°C or 32°C. Density was estimated from cell counts in 25-300μl samples under a dissecting microscope, taken in 6-10h intervals, for a total of 80h. Using FISH, we determined the proportion of infected cells after c. 48h, when populations were reaching carrying capacity.

Due to very low levels of infection or even entire loss of infection in the 32°C long-term selection treatment (see Results), all selection lines from strain 47 and one from strain 51 were omitted from the assay. Thus, in total, we assayed 23 selection lines and the four ancestral strains, with a total of 324 replicates (27 populations x 3 monoclonal lines x 2 infection status (infected / cured) x 2 assay temperatures).

### STATISTICAL ANALYSIS

We analysed variation in three parameters:

1. Infection persistence, i.e., the proportion of infected individuals in an (infected) assay replicate after 48h. This measure is related to the fidelity of vertical transmission during periods of host population growth as well as to the resistance of the parasite to the high temperature exposure (32°C).
2. Virulence, taken as the difference in growth between infected and cured assay replicates. Infection influenced both growth rate (r) and carrying capacity (K) (Fig. S2). To obtain a combined measure, we calculated the Area Under the Curve (AUC) of density for each assay replicate after 56h, using the trapezoidal method. The AUC represents the cumulative density during the assay, weighted for the time interval between measurements (Capaul and Ebert 2003; Adiba et al. 2010). The AUC is convenient, because it summarises growth in a single value per replicate. We calculated the AUC over the first 56h to remain close to the estimates of infection prevalence (≈48h), but nonetheless integrate values close to carrying capacity (correlation between AUC_56h_ and AUC_80h_: r = 0.96, n = 324, p < 0.0001). For each combination of monoclonal line and assay temperature, there was one infected and one cured assay replicate, and virulence was calculated as the difference between them: [(AUC_infected_) / (AUC_cured_)] −1. Thus, negative values indicate a negative effect of infection on host population growth. We did not specifically correct these estimates for loss of infection at 32°C assay temperature. There was no significant relationship between assay infection prevalence after 48h and virulence (r = 0.07, n = 81, p > 0.5), and adding prevalence as a covariate produced no significant changes in the main analysis of virulence (main effect of prevalence, interactions with other terms in the model: all p > 0.3).
3. Vertical transmission rate, calculated as: [(infection prevalence)_48h_ x (density)_48h_ - (density)_0h_] / [(density)_0h_ x 48h]. Integrating over host reproductive rate and the fidelity of vertical transmission, this measure describes the proliferation of infected cells and therefore represents an estimate of the vertical transmission rate (R_0_) of *Caedibacter* (Mangin et al. 1995).

Using General Linear Models (GLM), we analysed variation in infection prevalence with a binomial error structure (logit link), and variation in virulence and in (square-root-transformed) vertical transmission rate with a normal error structure. We fitted fully factorial models with selection temperature (long-term 26°C, 32°C or ancestor), assay temperature (26°C or 32°C) and strain genetic background (A30, 116, 51, 298) as model terms. Selection line and monoclonal line identity were included as nested terms and considered as random factors for hypothesis testing. Post-hoc contrasts compared selection treatments separately for each assay temperature and strain background.

We performed two analyses of trait relationships. First, we performed multiple regressions to test how changes in vertical transmission rate (response variable) depended on change in infection persistence and/or change in virulence (explanatory variables). Trait change was calculated as the difference between the value of each evolved line and the mean of the corresponding ancestral strain. All selection lines were pooled and analyses performed for 26°C and 32°C assay temperatures, respectively. Second, in a Covariance Analysis (Bell 1989; He et al. 2009; Pariaud et al. 2013), we tested whether experimental long-term treatments affected the covariance between vertical transmission rates at 26°C and at 32°C. This analysis considers data at 26 and 32°C as joint response variables. For example, a negative covariance would be indicative of a trade-off. Analogous to variance partitioning in univariate ANOVA, it allows covariance partitioning in multifactorial models (for details, see Bell 1989). In our case, model terms were long-term treatment, strain genetic background and selection line identity. Analyses were performed with JMP (SAS Institute 2017).

## Results

### DEMOGRAPHY AND EPIDEMIOLOGY DURING THE LONG-TERM SELECTION EXPERIMENT

Over the 52 weeks of the experiment, we observed a negative effect of high-temperature stress on *Paramecium* cell density (Fig. 1a). While density at 26°C reached levels of over 4000 cells per ml, it rarely exceeded 1000-2000 cells per ml at 32°C. Furthermore, infection remained at nearly 100% in the 26°C control lines, but generally declined at 32°C, with four selection lines (all from strain 47 and one from strain 51) even losing infection completely (Fig. 1b). Only for one strain (298), infection remained at nearly 100% at 32°C.

**Fig. 1.**
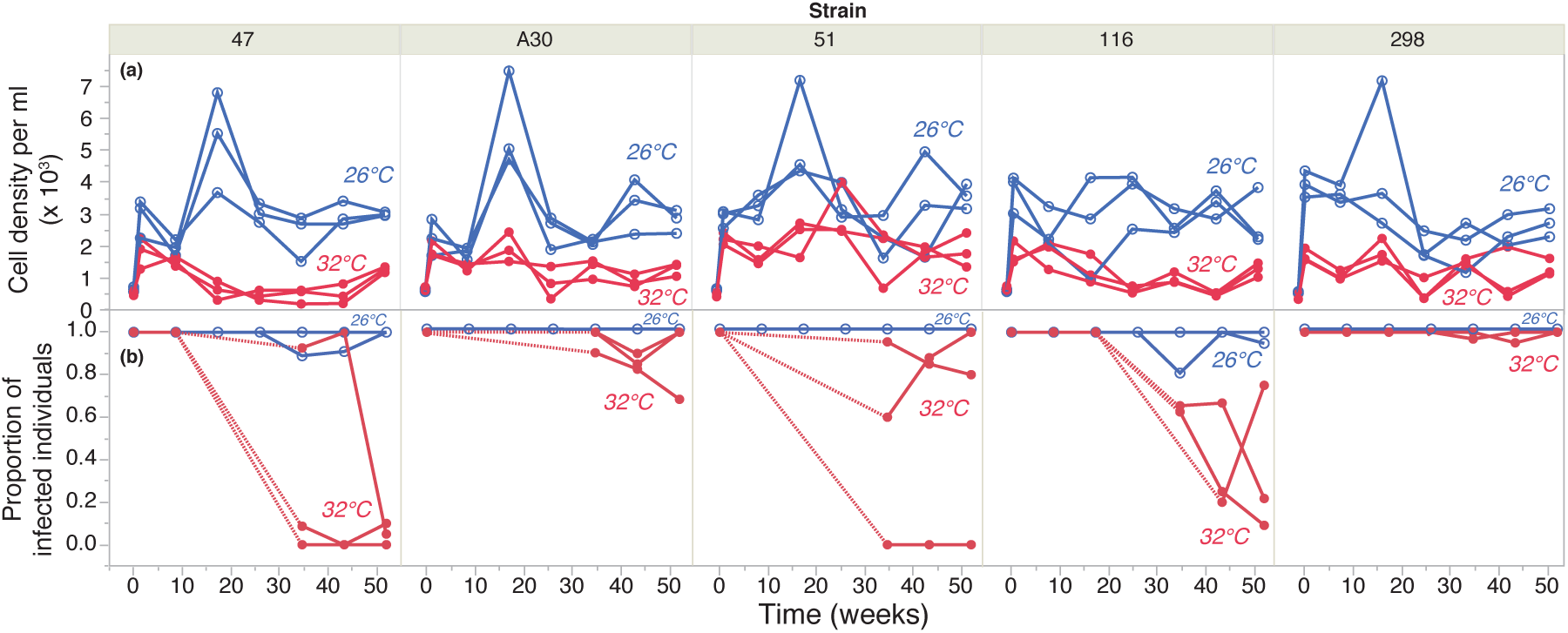
Population density (a) and infection prevalence (b) in individual long-term selection lines from each of five strain genetic backgrounds (47, A30, 51, 116, 298) propagated at 26°C (blue symbols) or 32°C (red symbols), over the course of 52 weeks (≈ 150 host generations). Densities were taken in 2-month intervals. Only pooled prevalence data (combined over selection lines from the same strain and treatment) were available for the first half of the experiment, and trajectories are shown as stippled to indicate uncertainty of divergence of selection lines (i.e., when pooled prevalences were < 1). After week 30, prevalences are shown for individual long-term lines.

### ADAPTATION ASSAY

#### Responses to selection

##### Infection persistence

Because the fit for the full statistical model did not converge (due to nearly no variation at 26°C assay temperature), we run models separately for each assay temperature (Table S1). At 26°C assay temperature, infection prevalence remained at 100% in more than 90% of the replicates (mean proportion infection: 0.99 ± 0.003), and the analysis did not detect any significant effects of genetic background or selection treatment (Fig. 2a, Table S1). In contrast, infection prevalence generally decreased over the 48h at 32°C (Fig. 2b). This decrease varied significantly with strain identity (F_3, 15_ = 10.47, p = 0.0006) and with selection treatment origin (F_2, 15_ = 12.53, p = 0.0006; Table S1). Namely, prevalence was less reduced in lines from the 32°C selection treatment (−20% or less prevalence reduction) than in ancestral lines or lines from the 26°C selection treatment (up to −60% reduction; Fig. 2b), indicating a general positive direct response to selection for infection persistence at this temperature.

**Fig. 2.**
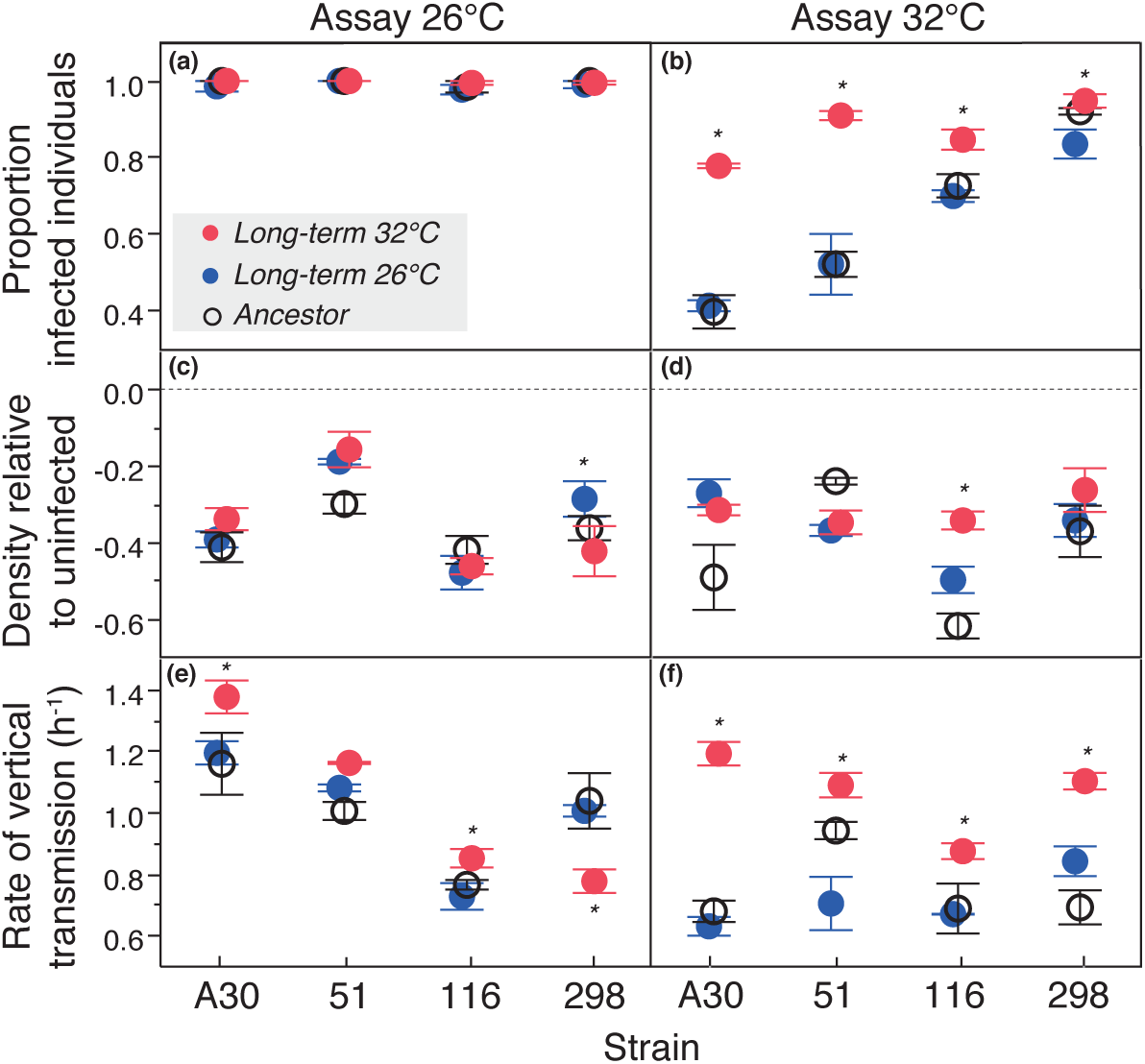
Infection prevalence (a, b), virulence (c, d) and vertical transmission rate (e, f) in the adaptation assay, carried out at 26°C and 32°C assay temperature. Each panel shows the mean (± SE) for selection lines from 26°C (blue symbols) or 32°C (red symbols) selection treatments as well as the corresponding ancestral lines (open symbols), from each of four genetic strain backgrounds. Asterisks indicate significant differences (p < 0.05) between 26°C and 32°C selection treatments, as revealed by post-hoc contrasts.

##### Virulence

We found substantial levels of virulence for all selection lines (Fig. 2c, d). On average, infected lines produced 40% lower densities than their cured, symbiont-free counterparts (t_23_ = 19.9, p< 0.0001). There was an overall trend of evolution towards lower virulence (mean difference evolved vs ancestral lines: t_22_ = 4.25, p = 0.0002). This virulence reduction in evolved lines tended to be expressed more clearly when lines were measured at 32°C (assay temperature x selection temperature: F_2, 15_ = 4.96, p = 0.0222; Table S2; Fig. 2d). Unlike for infection persistence, there was little evidence for specific adaptation to the long-term temperature treatments. Instead, detailed analysis revealed highly variable patterns for different strain genetic backgrounds (assay temperature x selection temperature x strain background: F_6, 15_ = 10.8, p = 0.0001; Table S2). Thus, in only two cases, ’local’ selection lines were less virulent than foreign lines (strain 298 evolved at 26°C, Fig. 2c; strain 116 evolved at 32°C, Fig. 2d). In the other cases, local and foreign selection lines showed relatively similar responses, or even increased rather than decreased virulence at 32°C relative to the ancestor (strain 51 evolved at 32°C; Fig. 2d). In other words, there was no clear general trend in the relationship between direct responses to selection in one temperature environment and correlated responses to selection in the other.

##### Vertical transmission rate

Realised rates of vertical transmission are a function of infection persistence and host replication rate (≈ infection prevalence x host density). Differences in vertical transmission between evolved and ancestral lines varied significantly with assay temperature (assay temperature x selection temperature: F_1, 15_ = 68.98, p < 0.0001; Table S2). At 32°C, there was a clear positive direct response to selection across all strain backgrounds (Fig. 2f): Lines selected at this temperature showed up to twice as high vertical transmission rates than ancestral lines or lines from the 26°C selection treatment. At 26°C, patterns varied with strain genetic background (assay temperature x selection temperature x strain background: F_6, 15_ = 5.35, p = 0.0039; Table S2), but in no case, selection at 26°C seemed to have consistently improved vertical transmission over that of ancestral backgrounds (Fig. 2e).

If anything, certain lines selected at 32°C showed the highest transmission rates, meaning that correlated responses to selection at 32°C were higher than the direct responses to selection at 26°C.

#### Trait correlations

##### Multiple regression analysis

Analysis of the relationships between trait changes (Table S3) showed that improvement in vertical transmission at 26°C (be it through direct or correlated responses to selection) was primarily associated with a decrease in virulence (F_1, 20_ = 10.58, p = 0.0040; Fig. 3). At 32°C, both increases in infection persistence (F_1, 20_ = 9.96, p = 0.0050) and decreases in virulence (F_1, 20_ = 4.42, p = 0.0483) made significant contributions to improved vertical transmission. However, the path through persistence change produced a stronger signal than that through virulence change (Fig. 3).

**Fig. 3.**
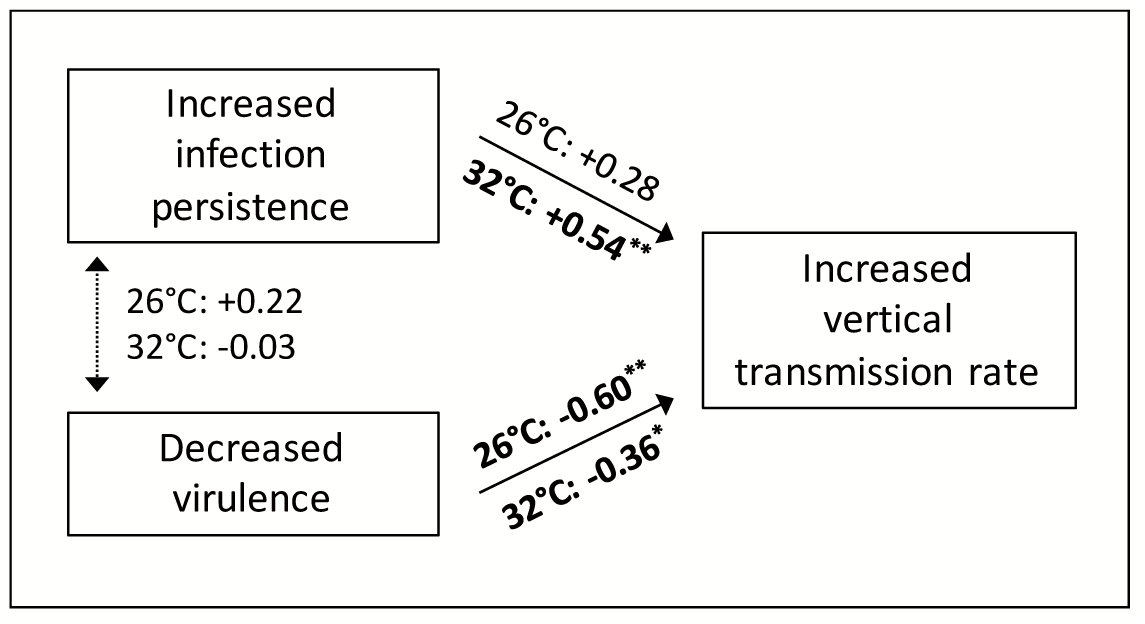
Relationships between trait changes, as revealed by multiple regression, using change in vertical transmission rate as response variable and changes in infection persistence and virulence as explanatory variables. Trait change was calculated as the difference between evolved and ancestral lines. Separate analyses were performed for 26°C and 32°C assay temperatures, respectively. Values for unidirectional arrows represent standardised beta regression coefficients; values for bidirectional arrows are correlation coefficients. For convenience, we denote the direction of change for each trait (increase or decrease relative to ancestor); the coefficients indicate the relationship between these increases / decreases.

##### Trade-off analysis

We found little evidence for trade-offs involved in temperature-specific adaptation. If anything, the correlation between vertical transmission rate at 26°C and 32°C was positive, rather than negative (all monoclonal lines combined, with two outliers removed: r = 0.19; n = 79, p = 0.0953). Although this correlation was only weakly positive, the covariance analysis (Table S4) revealed significant effects of strain genetic background (F_3, 15_ = 189, p < 0.0001), meaning that some genetic backgrounds have higher vertical transmission rates than others, at both 26 and 32°C. We also detected a significant effect of selection treatment (F_2, 15_ = 213.8, p < 0.0001): Namely, several selection lines selected at 32°C extended the trait range of the ancestral lines (grey circle in Fig. 4) and showed highest vertical transmission rates at both 26 and 32°C (points in top right corner in Fig. 4). This was the consequence of positive direct and correlated responses to selection at 32°C for these lines (see also Fig. 2e, f).

**Fig. 4.**
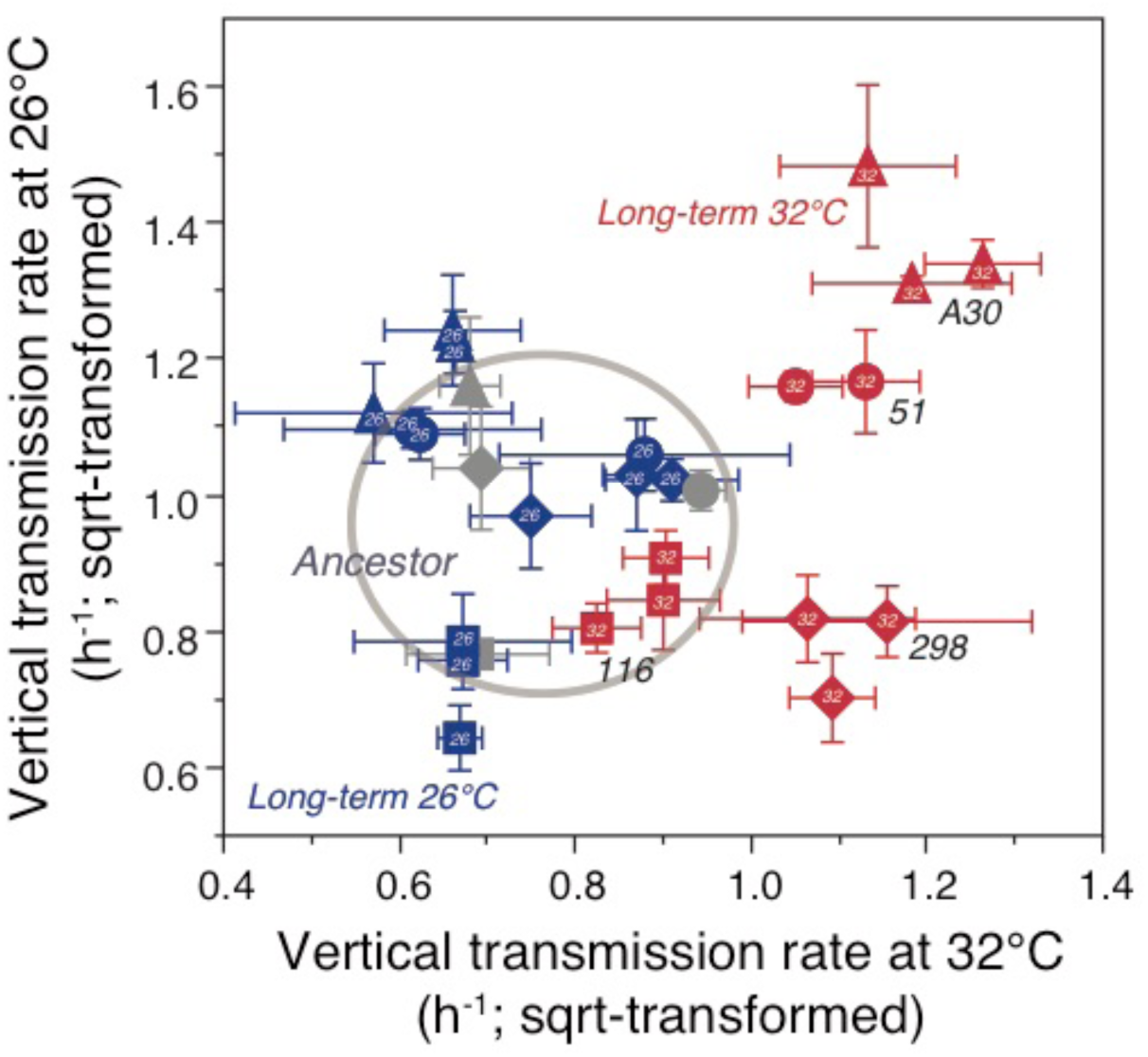
Relationship between vertical transmission rate at 26°C and at 32°C assay temperature, shown for selection lines evolved at 32°C (red) or 26°C (blue) and for ancestral lines (grey). Each point represents the mean (± SE) for an individual selection line. The grey circle illustrates the ancestral trait range. Different symbols refer to four different strain backgrounds (116, 298, 51, A30); the origin of the temperature selection treatment (26°C or 32°C) is shown within each symbol.

## Discussion

This study investigated how a host-symbiont system with obligate vertical transmission adapts to environmental stress. Because strict vertical transmission links host and symbiont fitness, we had hypothesised that adaptation to heat stress in our experimental *Paramecium-Caedibacter* system may involve the evolution of lower virulence or even benevolence, similar to symbiont-mediated heat-stress protection in aphids (Russell and Moran 2006; Douglas 1998). We found clear evidence for improved vertical transmission rates in long-term lines evolving under temperature stress (32°C). There also was a general, temperature-independent trend towards lower virulence, but in no case infection with *Caedibacter* became beneficial to the host.

### EVOLUTION OF INCREASED INFECTION PERSISTENCE AND VERTICAL TRANSMISSION

Previous work had demonstrated that *Caedibacter* is less heat-stress tolerant than its host (Dusi et al. 2014). Consistent with these results, ancestral strains and selection lines from the 26°C long-term treatment showed substantial reductions in infection prevalence when exposed to 32°C in the 48-h adaptation assay. This decline was also observed over the course of the 32°C long-term treatment, but infection nonetheless persisted in the majority of selection lines until the end of the experiment. These lines retained consistently higher infection prevalence and substantially higher rates of vertical transmission at this temperature in the assay.

A straightforward explanation for these observations is adaptation of *Caedibacter* to high-temperature stress, just like it is observed in experiments with free-living bacteria (Bennett et al. 1992; Saarinen et al. 2018), where adaptation of growth rate occurs over similar time spans. Increased heat resistance may therefore allow *Caedibacter* to attain viable within-host densities, track the regular dilution, when host cells divide, and thereby achieve a high fidelity and rate of vertical transmission. Adaptation was highly consistent over the four strain backgrounds tested, indicating a general evolutionary potential to respond to temperature stress.

It is often assumed that local adaptation to novel environments causes negative correlated responses to selection in other environments (Kassen 2002; Nidelet and Kaltz 2007). Here, however, selection for increased vertical transmission at 32°C did not seem to trade off with that at 26°C. In fact, the overall quasi-genetic correlation even tended to be positive, such that higher rates of vertical transmission at one temperature were associated with higher rates at the other (Fig. 4). The absence of fitness costs of adaptation to high-temperature stress at permissive temperatures (such as 26°C here) is not uncommon (Bennett and Lenski 2007; Killeen et al. 2017), and their detection perhaps requires fitness measurements at the lower end of the species’ specific temperature range (Killeen et al. 2017). Our analysis of the covariance between the two present temperature environments (26°C, 32°C) further revealed significant contributions from the selection treatment. Namely, the lines with the highest rates of vertical transmission at both temperatures were from 32°C selection treatment (Fig. 4, top right corner). Again, this finding resembles results for free-living bacteria, where temperature stress did not only select for an increased temperature tolerance, but also for general improvement in fitness (Bennett and Lenski 1993, 1996). Extrapolated to a (hypothetical) geographical context, our results would then suggest that infected *Paramecium* from the ’marginal’ 32°C environments can invade the permissive 26°C ’mainland’ (see also Killeen et al. 2017). Indeed, theoretical papers have highlighted the importance of selection in marginal habitats for range expansions, even though these investigations generally focus on the reverse question of how dispersal from the mainland affects adaptation at the margins (Kirkpatrick and Barton 1997; Sexton et al. 2009).

### VIRULENCE EVOLUTION

Theory predicts that virulent symbionts with exclusive vertical transmission can persist only under very limited conditions (Ewald 1987; Jones et al. 2007). One possibility is that fitness costs are compensated through protection against biotic or abiotic stress (Brownlie and Johnson 2009; Jones et al. 2011). *Caedibacter* infection causes constitutive over-expression of heat-shock proteins (Grosser et al. 2018), which suggested the potential for infection to even become beneficial to the host under temperature stress (e.g., Hori and Fujishima 2003). This was not the case here. In all evolved lines, infection still substantially decreased host division relative to uninfected lines.

Nonetheless, when compared to the ancestral lines, there was a general trend towards decreased virulence over all evolved lines combined. This can be explained by the protocol of our long-term experiment, where recurrent population dilution imposed a general selection pressure for increased division rate and, by extension, increased rates of vertical transmission (see Magalon et al. 2010). Indeed, our analyses show that evolutionary increases in vertical transmission were associated with decreased virulence in both selection environments.

Importantly, however, there was no clear general signal of temperature-specific virulence evolution. Instead, we found strong interactions between strain background, selection temperature and assay temperature. Thus, only lines from strain 116 seemed to have evolved specifically low virulence at 32°C, whereas in the other three cases, changes in virulence were independent of the selection treatment, or even showed the opposite trend of increased virulence (Fig. 2e, f). These variable outcomes explain why virulence evolution made a relatively weak contribution to vertical transmission evolution at 32°C (Fig. 3).

General theory holds that exclusively vertically transmitted parasites cannot be maintained if transmission is imperfect or if they reduce host fitness (Lipsitch et al. 1996). This verdict likely also holds for our *Caedibacter*-infected populations in the 32°C environments. Even though revealing clear signs of adaptation, the assay showed that vertical transmission is not perfect in the 32°C-adapted lines, causing the spin-off of a considerable number of uninfected cells in only 48h during the short-term adaptation assay. Repeating such periods of high population growth (compared to the relatively low growth rates in the long-term selection experiment) should therefore lead to a continuous decline in infection prevalence. These uninfected ’escapees’ cannot be reinfected through horizontal transmission and this explains why infection prevalences in the long-term populations did not return to 100%, despite the improved rates of vertical transmission (Fig. 1b).

### CAVEATS AND LIMITATIONS

Given the observed levels of virulence and incomplete vertical transmission in our assay, it is surprising that heat-adapted infected long-term lines have persisted until the end of the experiment. In fact, even if we assume improved vertical transmission at 32°C, uninfected individuals should gradually take over the populations in 50-60 generations (as found for strain 47, Fig. 1b). It is possible that virulence is overestimated in monoculture assays and that direct interactions in competition assays reveal more accurate (i.e., lower) values. This also points to a potentially important missing piece in the picture, namely the capacity of *Caedibacter*-infected hosts to eliminate uninfected *Paramecium* (Schrallhammer and Schweikert 2009). This killer trait may compensate even large costs of infection and prevent the spread of newly cured individuals. We have not measured killer activity at 32°C, but the persistence of uninfected cells at this temperature clearly indicates that it is impaired (see also Dusi et al. 2014), similar to high-temperature sensitivity of frequency distorter functions in the vertically transmitted *Wolbachia* bacteria (Hurst et al. 2000). Ancestral strains differ in killer activity at 26°C (E. Dusi, unpublished data), but these differences do not readily explain the observed variation in infection persistence in our present assays (not shown). This calls for more detailed investigations of temperature-dependent killer trait expression and its potential to evolve at 32°C.

In systems with strict uniparental (asexual) vertical transmission, host and symbiont genomes are locked up within the same line of descent, and interaction traits may therefore be particularly likely to evolve as “shared traits” (Restif and Koella 2003). Separation of host and parasite contributions to the observed results would require artificial infection of the parasite in ’common garden’ host backgrounds by micro-injection techniques, which are currently not available for our system. To assess possible host evolution (e.g., Killeen et al. 2017), we performed additional analyses comparing growth (AUC of cumulative density) between cured evolved and cured ancestral lines. This analysis revealed no sign of adaptation of the *Paramecium* to the 32°C treatment (overall evolved vs. ancestor: t_43_ = 0.61, n.s.; interaction strain background x evolved/ancestor: F_3,7_ = 1.95, p > 0.2), suggesting that increased parasite persistence and vertical transmission are best explained by parasite adaptation.

### IMPLICATIONS AND CONCLUSIONS

The robustness of “nested systems”, such as host-symbiont interactions, against environmental variation may critically depend on how the symbionts perceive the external environment. Temperature is a factor that can reach symbionts relatively unfiltered and is known to be a limiting factor of their survival and reproduction in uni- and multicellular hosts (Hood et al. 2010; Paaijmans et al. 2010; Duncan et al. 2011, 2017). Here we demonstrate the possibility of within-host adaptation of an obligate bacterial endosymbiont to external high-temperature stress, possibly by the same mechanisms as their free-living counterparts. Knowing the potential of symbionts and parasites to evolve such adaptations may critically determine our predictions regarding their future geographic range under changing climatic conditions (Harvell et al. 2002; Lafferty 2009). We found that adaptation to the “novel” environment was largely independent of strain genetic background and cost-free with respect to performance in the “original” temperature environment. Such universality may greatly facilitate the emergence and spread of newly acquired adaptations.

Strict vertical symbiont transmission is considered a perfect prerequisite for the evolution of mutualism (Ferdy and Godelle 2005; Moran et al. 2008; Leigh 2010), in particular when the symbiont provides protection against hostile environments or natural enemies (Oliver et al. 2003). Indeed, experimental limitation of the horizontal pathway in parasites with a mixed mode of transmission has been shown to lead to evolutionary shifts towards increased vertical transmission and “benevolence” towards their hosts (Bull et al. 1991; Magalon et al. 2010; Shapiro and Turner 2018). Our study is different from these experiments. First, we used a parasite with already naturally obligate vertical transmission and, second, we focused on the consequences of its adaptation to a particular environmental condition (temperature). We found little evidence for temperature-specific adaptation of virulence, and despite the evolution of increased rates of vertical transmission, there seemed to be a general limit, below which virulence could not be reduced. This may reflect a baseline level of host resources *Caedibacter* needs to ensure its own reproduction and successful vertical transmission. Thus, just like for parasites with horizontal transmission (van Baalen and Sabelis 1995), fitness of vertically transmitted parasites may be maximised for optimal levels of virulence.

## Acknowledgements

We thank Alison Duncan and Emanuel Fronhofer for comments and discussion. Finn Pond provided *Paramecium* strains. German Research Foundation (DFG priority program Host-Parasite Coevolution: BE-2299/5-1; RA-1920/1-1) and European FP7 program IRSES (247658) provided financial support.

## Data archiving

The database of the adaptation assay will be archived in the Dryad Digital Repository.

