## Supplementary Information for "When enemies do not become friends: Experimental evolution of heat-stress adaptation in a vertically transmitted parasite"

Supporting information

Table S1. Analysis of Deviance of infection persistence (= proportion of infected cells) after c. 48h in the adaptation assay at 26°C and 32°C, respectively. Generalised Linear Model, using logit link function and binomial error structure. Strain genetic background and long-term treatment (26°C, 32°C, ancestor) were the explanatory factors. Selection line was considered a random factor and used as denominator in quasi-F tests, based on mean deviances (MD).

| | | Assay 26°C | | | Assay 32°C | | |
|---|---|---|---|---|---|---|---|
| Source | d.f. | MD | F | p | MD | F | p |
| Strain background | 3 | <0.001 | <1 | n.s. | 30.67 | 10.47 | 0.0006 |
| Long-term treatment | 2 | <0.001 | <1 | n.s. | 36.72 | 12.53 | 0.0006 |
| Strain*treatment | 6 | <0.001 | <1 | n.s. | 2.11 | 0.72 | 0.6400 |
| Selection line(strain, treatment) | 15 | 0.888 | 2.86 | 0.0024 | 2.93 | 2.57 | 0.0307 |
| Residual | 54 | 0.310 | | | 1.14 | | |

Table S2. Analyses of Variance of virulence (= cumulative density of infected *Paramecium* relative to cured counterparts) and of vertical transmission rate (sqrt-transformed), over 56h in the adaptation assay. Explanatory factors were strain genetic background, long-term treatment (26°C, 32°C, ancestor) and assay temperature (26°C, 32°C); selection line and monoclonal identity were considered as nested random factors. The 'Denom.' column specifies the model terms used as denominators in F-tests. MS = mean squares.

| | | | Virulence | | | Vertical transmission rate | | |
|---|---|---|---|---|---|---|---|---|
| Source | Denom. | d.f. | MS ($x10^{-2}$) | F | p | MS ($x10^{-2}$) | F | p |
| Strain background | [1] | 3 | 22.1 | 11.50 | 0.0004 | 47.7 | 32.89 | <0.0001 |
| Long-term treatment | [1] | 2 | 4.4 | 2.29 | 0.1356 | 71.5 | 49.29 | <0.0001 |
| Strain*treatment | [1] | 6 | 1.7 | 0.89 | 0.5236 | 10.9 | 7.53 | 0.0007 |
| Assay temperature | [2] | 1 | 1.4 | 2.91 | 0.1084 | 91.2 | 68.98 | <0.0001 |
| Strain*assay temperature | [2] | 3 | 3.0 | 6.41 | 0.0052 | 23.4 | 17.72 | <0.0001 |
| Treatment*assay temperature | [2] | 2 | 2.3 | 4.96 | 0.0222 | 44.0 | 33.30 | <0.0001 |
| Strain*treatment*assay temperature | [2] | 6 | 5.0 | 10.80 | 0.0001 | 7.1 | 5.35 | 0.0039 |
| Selection line(strain, treatment) [1] | [3] | 15 | 1.9 | 2.38 | 0.0103 | 1.5 | 1.31 | 0.2288 |
| Selection line*assay temperature(strain, treatment) [2] | [4] | 15 | 0.5 | 0.64 | 0.8248 | 1.3 | 0.73 | 0.7395 |
| Monoclonal line(selection line, strain, treatment) [3] | [4] | 54 | 0.8 | 1.17 | 0.2870 | 1.7 | 0.93 | 0.6078 |
| Residual [4] | | 54 | 0.7 | | | 1.8 | | |

Table S3. Multiple regression analyses of response to selection in vertical transmission rate (sqrt-transformed), as a function of the responses to selection (RS) in infection persistence and in virulence. Analyses carried out separately for each assay temperature (26°C, 32°C). Responses to selection were calculated as differences in traits between evolved and ancestral lines.

| | | Assay 26°C | | | Assay 32°C | | |
|---|---|---|---|---|---|---|---|
| Source | d.f. | MS ($x10^{-2}$) | F | p | MS ($x10^{-2}$) | F | p |
| RS infection persistence | 1 | 3.6 | 2.27 | 0.1478 | 37.3 | 9.96 | 0.0050 |
| RS virulence | 1 | 16.6 | 10.58 | 0.0040 | 16.6 | 4.42 | 0.0483 |
| Residual | 20 | 1.6 | | | 3.7 | | |

Table S4. Analysis of the covariance between vertical transmission rate at 26°C and at 32°C (both sqrt-transformed), as a function of strain genetic background and long-term treatment (26°C, 32°C, ancestor). Selection line identity considered as a nested random factor. Analogous to mean squares, mean products (MP) from the covariance analysis were used for F-tests. The 'Denom.' column specifies the model terms used as denominators in these F-tests.

| Source | Denom. | d.f. | MP ($x10^{-2}$) | F | p |
|---|---|---|---|---|---|
| Strain background | [1] | 3 | 12.2 | 189.0 | <0.0001 |
| Long-term treatment | [1] | 2 | 13.7 | 213.8 | <0.0001 |
| Strain*treatment | [1] | 6 | 1.92 | 29.9 | <0.0001 |
| Selection line(strain, treatment) [1] | [2] | 15 | 0.06 | 1 | 0.4782 |
| Residual [2] | | 54 | -0.06 | | |

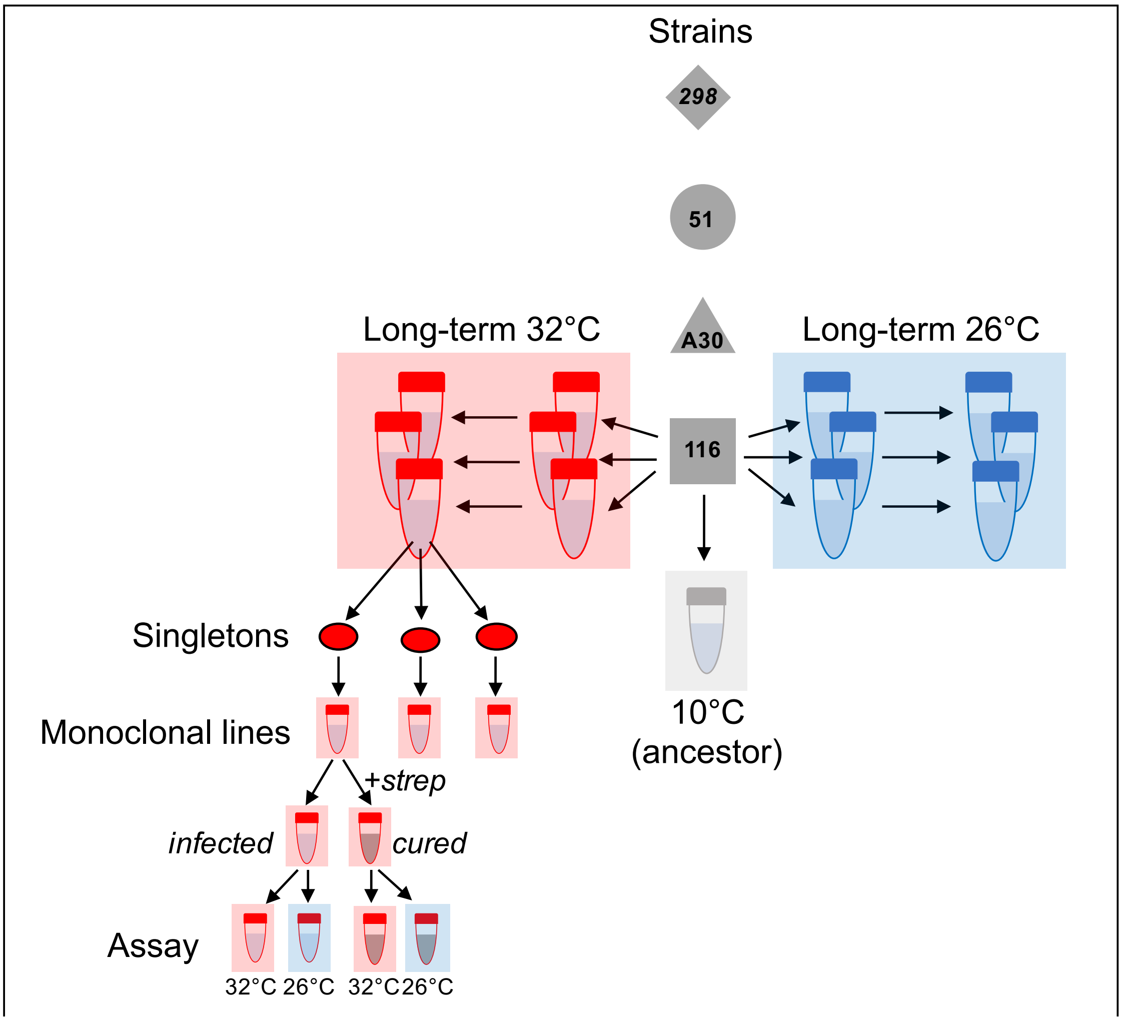


Fig. S1. Design of long-term selection experiment and set-up of adaptation assay. For each strain (= association of a given *Paramecium tetraurelia* strain with its own *Caedibacter taeniospiralis* symbiont), three long-term replicate selection lines were established at 26°C and 32°C, respectively. Selection lines were set up for five strains, but only the four strains indicated in the figure were used in the adaptation assay (strain 47 was not included in the adaptation assay due to quasi-total loss of infection; see main text). To preserve the ancestral state, additional lines were kept at 10°C. After c. 150 asexual generations, three infected single cells were isolated from each long-term line. These singletons were grown up as individual cultures, referred to as monoclonal lines. Monoclonal lines were then split and one half was cured from infection with streptomycin. In the assay, infected and cured monoclonal lines were then tested at 26°C and 32°C. The figure illustrates the setting up of long-term lines for one (116) of the four strains, and the preparation of monoclonal lines for one long-term selection line (selected at 32°C). Of course, the same protocols were used for all other strains and selection lines.

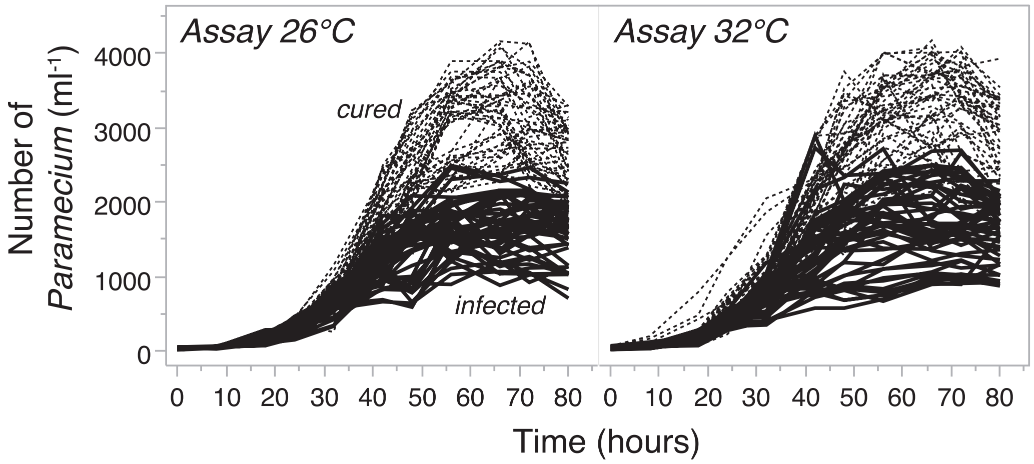


Fig. S2. Densities of infected and uninfected (= cured) replicates in the adaptation assay, measured at 26°C and 32°C assay temperature. Replicates represented small monoclonal cultures, which had been derived from the selection lines in the long-term experiment (see Fig. S1). The vertical lines indicate the end point (56h) chosen for calculations of the Area Under the Curve (AUC) of cumulated cell density. Differences in the AUC over this time period reflect variation in intrinsic growth rate and carrying capacity. The AUC was restricted to this period to be in concordance with the estimates of infection prevalence (≈48h).
